# Spies of the deep: an animal-borne active sonar and bioluminescence tag to characterise mesopelagic prey size and behaviour in distinct oceanographic domains

**DOI:** 10.1101/2023.10.19.563065

**Authors:** Mathilde Chevallay, Tiphaine Jeanniard du Dot, Pauline Goulet, Nadège Fonvieille, Cassandra Craig, Baptiste Picard, Christophe Guinet

**Author notes:** Corresponding author: Mathilde Chevallay, 405 route de Prissé-la-Charrière 79360 Villiers-en-Bois.

## Abstract

Mesopelagic fish, a central component of marine trophic networks, play a fundamental role in marine ecosystems. However, as they live in highly inaccessible environments, little information is currently available on their ecology, especially on the influence of oceanographic parameters on their distribution. The emergence of biologging technologies has made it possible to use deep-diving predators as bio-samplers of their environment in under-sampled regions. In this work, we deployed an innovative miniaturised sonar tag that combines active acoustics with high-resolution GPS, pressure, movement and light sensors on Southern elephant seals, a deep-diving predator feeding on mesopelagic prey. Seals were also equipped with oceanographic tags, allowing us to explore the functional relationships between oceanographic parameters, distribution and ecology of mesopelagic prey targeted by seals and the seals’ foraging behaviour. We highlighted strong vertical differences in prey characteristics and behaviour, with larger, more evasive and less bioluminescent prey in deeper waters. Moreover, prey encountered in warmer waters were found deeper, were more evasive and displayed a more marked diel vertical migration behaviour compared to prey encountered in colder waters, suggesting that prey accessibility and characteristics differ according to oceanographic domains. This study highlights the usefulness of the sonar-bioluminescence tag to infer mesopelagic prey distribution and habitat when deployed on deep-diving predators such as elephant seals.

## 1. Introduction

Mesopelagic fish represent the largest vertebrate biomass on Earth (Hidalgo and Browman, 2019; Irigoien et al., 2014; Proud et al., 2019) and play a fundamental role in marine ecosystems. They are a central component in the marine trophic network, allowing energy transfers between low trophic levels, i.e. phyto and zooplankton, and upper trophic levels, i.e. predators (Cherel et al., 2010). Most of mesopelagic fish perform diel vertical migrations, moving up to the surface at night and going back to deep waters during the day (Catul et al., 2011), and thus play a key role in the carbon pump by facilitating carbon transfer from the surface to deeper ecosystems (St. John et al., 2016). However, despite their ecological importance, the ecology of mesopelagic micronekton remains poorly known, particularly because they live in highly inaccessible environments. From the perspective of predicting the effects of climate change on marine ecosystems, there is a need to increase our knowledge on the influence of environmental parameters on their distribution.

Usual methods to study mesopelagic communities include net sampling and acoustic surveys. Net sampling consists of deploying a net from an oceanographic or fishing vessel in order to collect part of the individuals present in the area with the aim of making an inventory of the species, estimating size of individuals and their biomass. However, net avoidance is commonly observed in pelagic fish, leading to a biased in the species collected (Irigoien et al., 2014; Kaartvedt et al., 2012). Moreover, numerous trawls at different depths are necessary to obtain detailed information on the distribution of species, which may be challenging. In contrast, acoustic surveys describe the fine-scale distribution of organisms in the water column. Most oceanographic vessels are equipped with on-board echosounders that emit successive sound pulses reflected off any object they encounter in the water column (Misund, 1997). The echosounder then receives the returning echo and, depending on the return time, the distance between the echosounder and the object can be estimated. Active acoustics thus provides detailed information on the distribution of organisms but with limited effect on their behaviour. As the distance an acoustic wave can travel in water is inversely proportional to the frequency of that wave, echosounders must emit low frequency waves to sample a large part of the water column. However, low-frequency acoustic waves do not provide detailed echoes of the insonified objects (Misund, 1997). This limits the usefulness of echosounders to obtain fine-scale data on organisms in deeper parts of the water column. Moreover, both acoustic surveys and net samplings require expensive, time-consuming and logistically demanding oceanographic campaigns, that can be highly challenging to set up in remote areas such as polar regions, limiting spatial and temporal coverage.

Recent developments of biologging technologies, i.e. devices attached on animals to record information about their behaviour and ecology (Rutz and Hays, 2009), have largely increased our understanding of the ecology of marine predators and their environment (e.g. (Guinet et al., 2014, 2001; Slip et al., 1994; Yoshino et al., 2020)). By integrating high-resolution oceanographic sensors such as temperature, salinity, light or chlorophyll a sensors in tags that can be attached on large deep-diving predators, these predators can be used as bio-samplers of their environment (Fedak, 2013; Harcourt et al., 2019; Treasure et al., 2017), making it possible to collect environmental data in under-sampled regions. Investigating the foraging ecology, i.e. at sea distribution, diving behaviour, diet, of marine predators do also provide information on their cryptic and generally poorly known prey (Cherel et al., 2008). In that context and among deep-diving predators, Southern elephant seals (*Mirounga leonina,* Linneaus 1758, SES hereafter) are of special interest. SES breed on sub-Antarctic islands and alternate at-sea periods to feed and on-land periods to breed and moult. During their months-long foraging trips, SES travel great distances, up to thousands of kilometres, and perform continuous dives at depths ranging generally from 200 to 1000 m, and up to 2000 m. Therefore, by covering large horizontal and vertical ranges, SES are an ideal species to sample environmental conditions in these remote areas (e.g. (Bailleul et al., 2007; Charrassin et al., 2010; Dragon et al., 2010; Guinet et al., 2014; Tournier et al., 2021). They became a major component in the observation of the Southern Ocean oceanographic conditions (Fedak, 2013). While adult males forage mostly on benthic and benthopelagic prey such as squids, young males and females mainly target mesopelagic prey such as myctophids (Cherel et al., 2008).

Deploying high-resolution movement sensors such as accelerometers in addition to oceanographic and GPS sensors on deep-diving predators allows for identifying prey capture attempts (Suzuki et al., 2009; Viviant et al., 2010; Ydesen et al., 2014), providing valuable information on horizontal and vertical distribution of mesopelagic prey targeted by these predators (Guinet et al., 2014). However, fine-scale information on prey, such as their size or behaviour remain challenging to obtain. Miniaturised video cameras can be attached to marine predators foraging in clear and shallow waters (Goldbogen et al. (2017) on blue whales *Balaenoptera musculus*, Heithaus et al. (2002) on tiger sharks *Galeocerdo cuvier*, Papastamatiou et al. (2018) on white-tip sharks *Carcharhinus longimanus*, Semmens et al. (2019) on white sharks *Carcharodon carcharias*). However, SES often forage in the aphotic zone, which requires a light-source to illuminate the scene and capture exploitable images, leading to an increased power consumption of the tag. In addition, using a light source in these deep waters can significantly alter both predator and prey behaviours (Geoffroy et al., 2021; Rooper et al., 2015; Ryer et al., 2009), limiting the usefulness of video cameras to collect meaningful data about SES’ prey characteristics and behaviour. Furthermore, recording duration is limited to a few hours, over foraging which can last several months, providing short, but important, glimpse on the prey and foraging behaviour of those species. Very recently, an innovative miniaturised sonar tag has enabled a step forward in the study of fine-scale biotic environment in cryptic marine predators. This state-of-the-art tag, inspired by a toothed whale echolocation system, uses a high-click rate narrow, forward directed beam that can insonify prey at a distance up to 6 m in front of the animal. Data recorded by this tag provide information on prey aggregation type, escape behaviour and acoustic size without altering either predator or prey behaviour (Goulet et al., 2019). A more recent version of the sonar tag also includes a light sensor specifically developed to study bioluminescent behaviour of prey (Goulet et al., 2020), offering a unique opportunity to accurately describe behaviour and characteristics of prey targeted by cryptic marine predators (Chevallay et al., in revision).

In this context, the aim of this study was to provide a better understanding of the distribution and habitat of mesopelagic prey in the Southern Ocean. Using an innovative method to classify functional oceanographic domains (FOD hereafter) according to their temperature and salinity profiles recorded by conductivi ty-temperature-depth (CTD) tags (Fonvieille et al., in review; Pauthenet et al., 2017; Tournier et al., 2021) deployed simultaneously with sonar tags on female SES, we aimed at exploring the functional relationships between oceanographic parameters, distribution and ecology of mesopelagic prey targeted by seals and the seals’ foraging behaviour. Previous studies showed that seals visit different areas with contrasted oceanographic conditions during their foraging trips (Bailleul et al., 2007; Biuw et al., 2007) and that seals targeting distinct FOD differed in their foraging efficiency (Guinet et al., 2014; Richard et al., 2016). Therefore, we predicted that these contrasted habitats would be home to prey with diverse characteristics and behaviour, and that some habitats favour prey accessibility and/or prey quality (i.e. larger or less evasive prey). Specifically, we hypothesised that prey tend to be more evasive in warmer FOD but also less accessible, as suggested by Guinet et al. (2014) and Richard et al. (2016).

## 2. Material and methods

### 2.1. Device deployments and data collection

Data were collected on 9 post-breeding female Southern elephant seals in October 2018 (n = 3), October 2019 (n = 3) and October 2020 (n = 3), in Kerguelen Islands (49°20’S – 70°26’E). Females were captured with a canvas head-bag, anaesthetized with a 1:1 combination of Tiletamine and Zolazepam (Zoletil 100 – 0.7 to 0.8 ml/100 kg) either injected intravenously (McMahon et al., 2000) or using a deported intramuscular injection system. Females were measured to the nearest centimetre and weighted to the nearest kilogram. They were equipped with a neck-mounted Argos tag (SPOT-293 Wildlife Computers, 72×54×24 mm, 119 g in air), a back-mounted CTD tag (SMRU-SRDL, 115×100×40 mm, 680 g in air) and either a head-mounted DTAG-4 sonar tag (n = 7, 95×55×37 mm, 200 g in air, see Goulet et al. (2019) for further details on the device) or a DTAG-4 sonar-light tag (n = 2, 92×69×37 mm, 250 g in air). Tags were glued to the fur using quick-setting epoxy glue (Araldite AW 2101, Ciba). CTD tags recorded conductivity, temperature and depth at 0.5 Hz. Sonar and sonar-light tags were programmed to sample GPS position (up to every minute at surface), tri-axial acceleration (200 Hz), tri-axial magnetometer (50 Hz) and pressure (50 Hz). Sonar-light tags also integrated a light sensor specifically designed to sample bioluminescence events with a sampling frequency of 50 Hz (Goulet et al., 2020). The active sonar within the tags recorded acoustic backscatter returning from 10 µs pings with a centre frequency of 1.5 MHz, at a 25 Hz ping rate for 2018 and 2019 deployments and 12.5 Hz for 2020 deployments. The active sonar operated with a 3.4° aperture beam width and a 6 m range (Goulet et al., 2019). Sonar and sonar-light tags were set to record data one day out of two to increase recording duration. Tags were recovered in late December to January when females returned to shore to moult using the same capture and sedation methods. All experiments were conducted under the ethical regulation approval of the French Ethical Committee for Animal Experimentations (2019070612143224 (#21375)) and the Committee for Polar Environment.

### 2.2. Foraging behaviour

Sonar, movement and location data recorded by the sonar tags were analysed using custom-written codes and functions from www.animaltags.org in MATLAB version 2022b (The MathWorks, 2022). Statistical analyses were conducted in R software version 3.5.1 (R Core Team, 2018). GPS tracks were plotted in a map using QGIS version 3.10 (QGIS Development Team, 2019). Diving behaviour was assessed by using pressure data recorded by the sonar tags. Seals were considered as diving when depth exceeded 20 m.

### 2.3. Prey capture attempt identification

Prey capture attempts (PrCAs hereafter) were detected from the 200 Hz tri-axial acceleration data recorded by the sonar tags, by computing the norm of the differential of the tri-axial acceleration (norm-jerk hereafter), as described in Goulet et al. (submitted). Spikes in the norm-jerk higher than 400 m.s^-2^ were classified as PrCAs (Goulet et al., submitted). As prey may be encountered in patches or may elude capture, leading to a bout structure in prey strikes, strikes occurring less than 15 s from the previous strike were grouped in the same bout as described by Goulet et al. (submitted).

### 2.4. Functional oceanographic domain identification

Functional oceanographic domains (FOD hereafter) were classified according to their similarities in their temperature and salinity profiles, following the method described in (Fonvieille et al., in review). We first generated temperature and salinity profiles (TS) on a constant vertical grid from 20 to 200 m for each dive using temperature and salinity data collected by the CTD tags. This depth range was chosen to maximise temporal coverage especially at night when seals perform shallower dives (Tournier et al., 2021). We then fitted B-splines to the profiles with 20 coefficients estimated by polynomial regression (R package “fda”, (Ramsay and Silverman, 2005)). A bivariate functional principal component analysis (FPCA, R package “fda”) was applied on the resulting coefficients of the decomposition to extract principal modes of variability in the curves shapes (see Pauthenet et al. (2017) and Nerini et al. (2022) for more details). Finally, a model-based clustering (Bouveyron et al., 2019) was applied on the first principal components of the FPCA to gather TS profiles sharing similar thermohaline vertical structure (R package “mclust”, (Scrucca et al., 2016)). The appropriate number of clusters was defined using the Bayesian Information Criterion (Schwarz, 1978) and the Integrated Completed Likelihood (Biernacki et al., 2000), as described in Fonvieille et al. (in review). The resulting clusters was considered as representing different FOD and were plotted along the track of SES.

### 2.5. Prey characteristics and behaviour

#### 2.5.1. Sonar data analysis

Sonar data recorded during bouts were displayed as echograms, showing the time on the horizontal axis and the distance from the sonar tag on the vertical axis, extending from 10 s before the bout start time to 2 s after the bout end time (Goulet et al., submitted). Because of the large number of bouts detected (ranging from 4595 to 12039 per animal), only a random subsample of 5 to 10% of bouts performed during acoustic data recordings were analysed. Different variables describing predator-prey interactions were extracted manually from echograms following the method described by Goulet et al. (submitted): (1) number of prey, (2) prey escape behaviour, (3) prey acoustic size. (1) The number of prey was defined as the maximum number of independent echo traces within a same ping (Jones et al., 2008). It was scored as one, two or more than two, i.e. a school of prey. (2) Prey escape behaviour was identified from the closing speed between predator and prey, which will vary in case of prey escape event, resulting in a change in the slope of the prey echo trace (Goulet et al., 2019; Vance et al., 2021). (3) Prey acoustic size was estimated from the −20 dB echo pulse width measured on the widest part of the prey trace on evasive prey only to get an acoustic size estimate as close as possible to the prey length (Burwen et al., 2003).

#### 2.5.2. Bioluminescence data analysis

Bioluminescent flashes were defined as short high intensity spikes in the 50 Hz light data recorded by the sonar-light tags decimated at a 5 Hz sampling rate. Spikes in the data higher than 0.07 (arbitrary unit) were classified as flashes (Goulet et al., 2020). Flashes detected within less than 5 s of each other were considered coming from the same bioluminescence event (Goulet et al., 2020). If a bioluminescence event started between 5 s before and 5 s after the bout, it was considered associated with the bout. Bioluminescence events may consist of a series of short flashes, so the 5 Hz data may not have sufficient resolution to accurately describe them. Therefore, we used 50 Hz light data recorded during bioluminescence events to calculate the number of peaks in the light signal as well as the intensity and duration of each peak.

### 2.6. Statistical analyses

For the following analyses, only data recorded on seals equipped with sonar-light tags were used for the comparisons between flashing and non-flashing prey, while comparisons between evasive and non-evasive prey were made using data recorded by all the sonar-tagged seals. For all subsequent models, assumptions were checked and any deviation from them were corrected in the models.

#### 2.6.1. Inter-functional oceanographic domain differences in prey characteristics and distribution

Prey accessibility, using the depth of bout as its proxy (Boyd et al., 2015; Jouma’a et al., 2016) was compared between FOD using LMM (R package nlme, (Pinheiro et al., 2017)) with dive depth as response variable, FOD as fixed effect and seals’ individual identities as a random effect during both day and night. To test for differences in prey behaviour according to FOD, the proportion of evasive versus non-evasive prey and flashing versus non-flashing prey was compared between FOD using generalised linear mixed models with binomial distribution (GLMM hereafter, R package lme4, (Bates et al., 2015)). We used the proportion of each prey type as a response variable, FOD as fixed effect and seals’ individual identities as a random effect. As the two seals equipped with sonar-light tag almost never visited FOD4, we did not compute the proportion of flashing prey in this FOD. Prey flight initiation distances (m) were compared between FOD using LMM with flight initiation distance as response variable, FOD as fixed effect and seals’ individual identifies as a random effect. Bout duration (s), a proxy for prey capture difficulty, or an indication of multi-prey capture (Goulet, 2020), was compared between FOD using LMM with bout duration as a response variable, FOD as fixed effect and seals’ individual identities as a random effect. Finally, distribution of prey acoustic size was reconstructed for each FOD and were compared using a pairwise Kolmogorov-Smirnov test with Bonferroni correction for multiple testing.

#### 2.6.2. Intra-functional oceanographic domain differences in prey characteristics and distribution

Inside each FOD, difference in depths at which prey were encountered between evasive and non-evasive prey and between flashing and non-flashing prey were compared using prey encounter depth as response variable, prey type (evasive vs. non-evasive prey, flashing vs. non-flashing prey) as fixed effect and seals’ individual identities as a random effect. To test the hypothesis that prey size varies with depth, prey acoustic size distributions were reconstructed for depth classes every 200 m, from 20 to 1000 m, and were compared using Kolmogorov-Smirnov test with Bonferroni correction for multiple testing inside each FOD. Finally, the relationship between bout duration and bout depth was assessed for each FOD using a LMM with bout duration as response variable, bout depth as fixed effect and seals’ individual identifies as a random effect.

## 3. Results

### 3.1. Foraging behaviour and targeted prey

Tags recorded data during 29-79 days (Table 1). One GPS sensor and one CTD tag failed to record any data on individual ml18_293a and individual ml19_296a respectively so data recorded from these individuals were not used for spatial analyses. Over that period, seals performed 69 ± 7 (mean ± sd) dives per day (range 57-79 dives per day), and 511 ± 53 bouts/day (range 443-610 bouts/day). Seals performed dives 418 ± 93 m deep for 19 ± 3 min. A diel diving pattern was observed for almost all individuals, with deeper and longer dives performed during the day (456 ± 96 m, 21 ± 4 min) than during the night (331 ± 75 m, 16 ± 2 min, LMM, P < 0.001). Seals almost exclusively targeted single prey (99% of the targeted prey) while schools of prey represented only 1% of the targeted prey. Among single prey, 81% were evasive. For the two individuals equipped with sonar-light tags, flashing prey represented 24 and 29% of their targeted prey respectively. Prey acoustic size mainly distributed from 5.1 to 9.0 cm (1^st^ and 3^rd^ quartiles, Q1-Q3 hereafter), with a range of 1.2-32.8 cm.

**Table 1:** Summary of field deployments: length (cm) and weight (kg), number of days of recordings, number of recorded dives and bouts of female SES from Kerguelen Islands equipped with either sonar or sonar-light tag during their post-breeding foraging trip. Seal ID comprises the two first letters of the Latin species binomial followed by the year and Julian day of attachment, and a letter denoting the sequential animal of day.

| Seal ID | Weight<br>(kg) | Length<br>(cm) | Device | Number of<br>days of<br>recordings | Number of<br>recorded<br>dives | Number of<br>recorded<br>bouts |
| --- | --- | --- | --- | --- | --- | --- |
| ml18_292a | 237.5 | 226 | sonar | 43 | 3393 | 22223 |
| ml18_293a | 158.5 | 225 | sonar | 29 | 1760 | 6823 |
| ml18_294a | 249.5 | 226 | sonar | 35 | 2380 | 11694 |
| ml19_295a | 287.5 | 239 | sonar | 30 | 2096 | 14284 |
| ml19_295c | 321.5 | 237 | sonar | 35 | 2336 | 18885 |
| ml19_296a | 252.2 | 225 | sonar | 28 | 1752 | 8205 |
| ml20_295a | 380.0 | 268 | sonar-light | 77 | 7669 | 41439 |
| ml20_296a | 380.0 | 231 | sonar-light | 79 | 4535 | 40304 |
| ml20_296c | 380.0 | 196 | sonar | 73 | 5357 | 36684 |

### 3.2. Foraging area

All seals but two travelled east of the Kerguelen Islands (Figure 1). The two other seals travelled west. They concentrated their foraging activity at a mean distance from the shore of 1170 ± 539 km (Q1-Q3: 789-1663 km, Figure 1).

**Figure 1:**
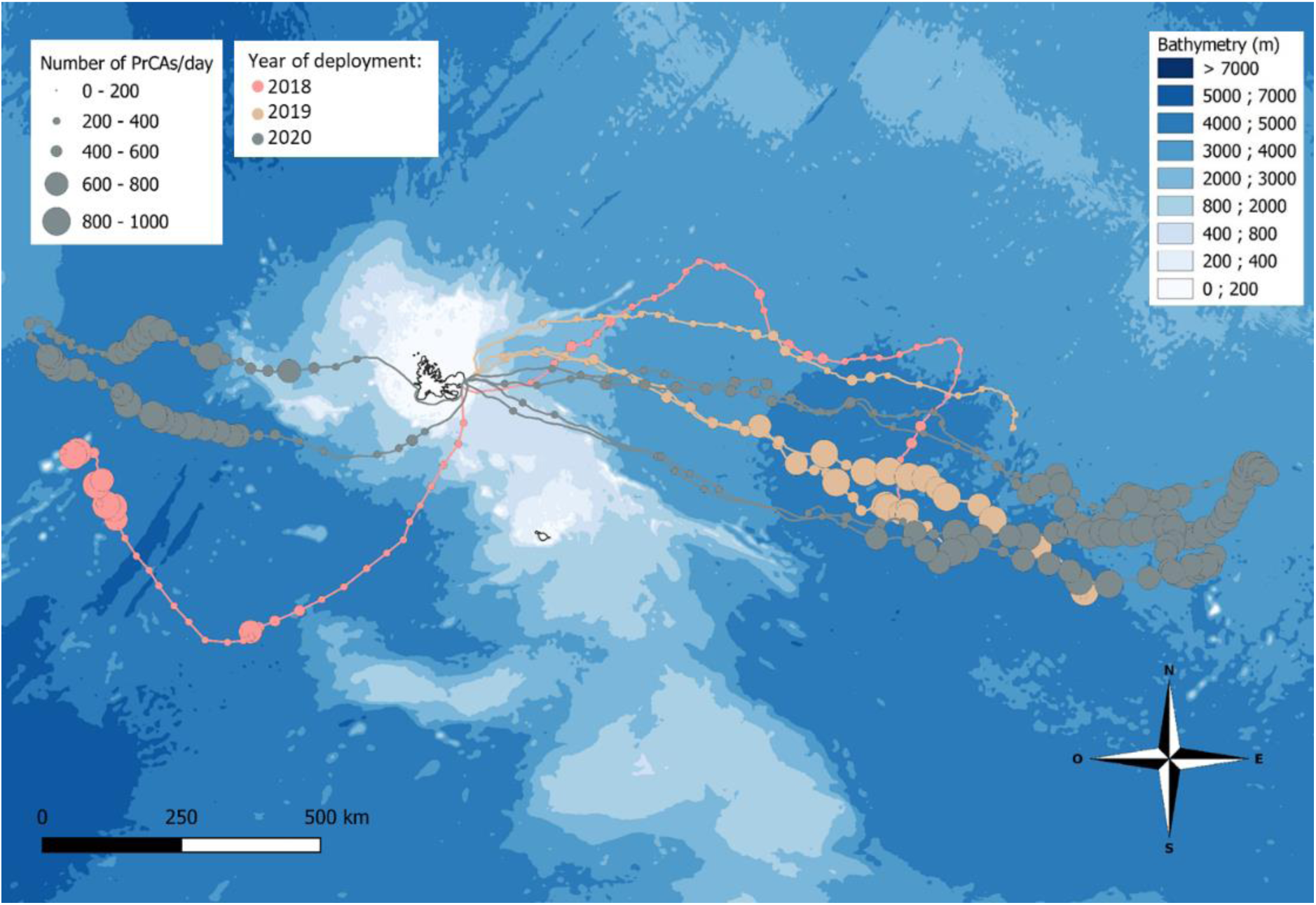
GPS tracks of eight female SES from Kerguelen Islands equipped with high-resolution sonar and movement tags in 2018 (n = 2, in pink), 2019 (n = 3, in beige) and 2020 (n = 3, in grey) during their post-breeding foraging trips (November-January). Each point represents one day, with the size of the point being proportional to the number of bouts performed that day. For 6 females, the tags stopped recording before the end of the foraging trip.

### 3.3. Inter-functional oceanographic domain differences in prey characteristics and distribution

During their foraging trips, seals encountered four FOD that we identified from *in-situ* temperature and salinity profiles recorded by CTD tags (Figure 2). These FOD were distributed along an increasing temperature gradient from FOD 1 to FOD 4, with little variation in salinity (Figure 2). Seals foraging East of Kerguelen visited mainly FOD 2 (46% of dives) and in comparison performed little feeding activity in the warmer FOD 1, 3 and 4 (22%, 22% and 11% of dives respectively, Table 2). In contrast, the two individuals foraging West of Kerguelen mostly concentrated their foraging activity in FOD 1 (79% of dives), the coldest FOD. They performed 21% of dives in FOD 2, and never visited FOD 3 and 4 (Table 2).

**Figure 2:**
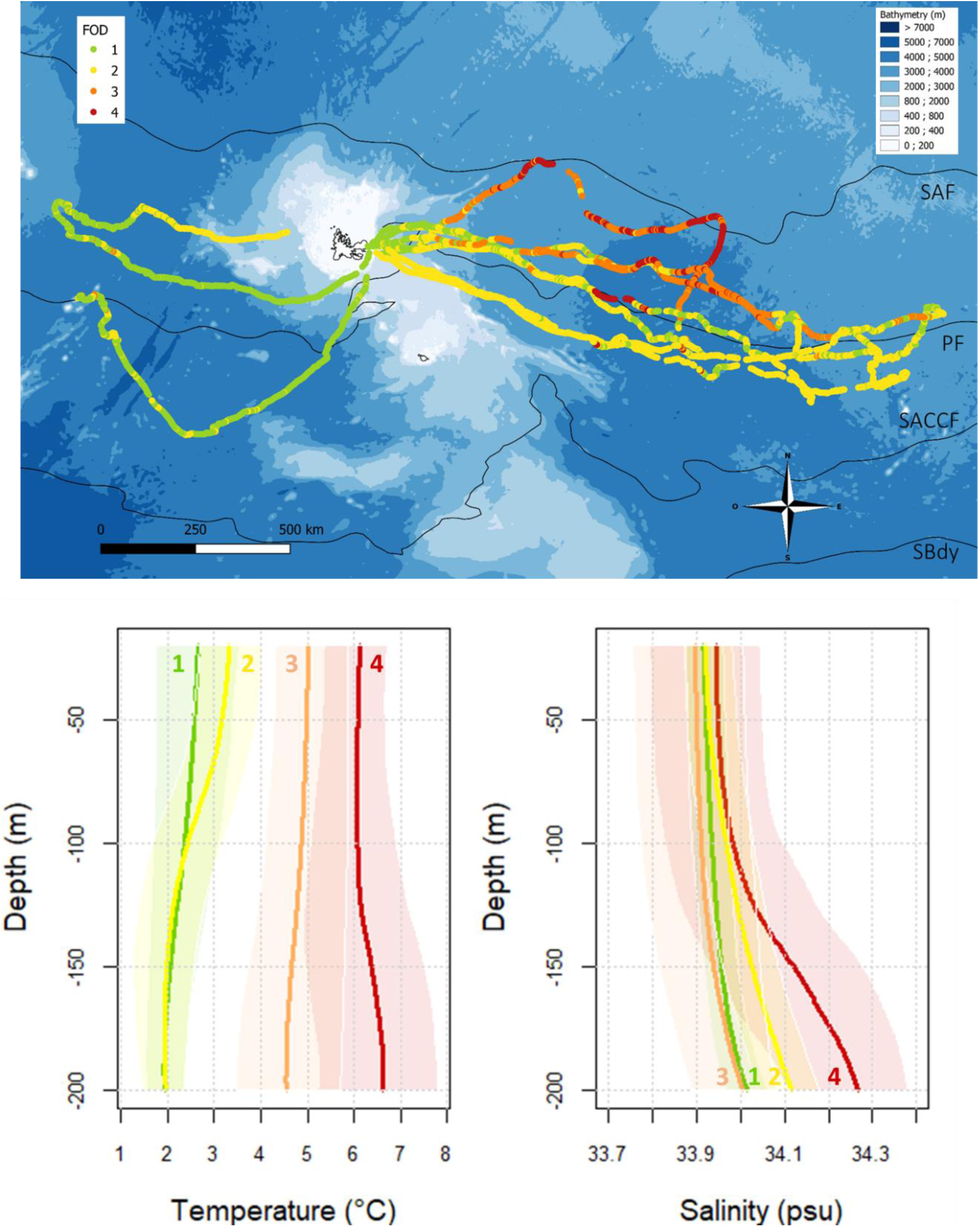
(A) GPS tracks of seven female Southern SES from Kerguelen Islands equipped with high-resolution sonar and movement tags in 2018 (n = 2), 2019 (n = 2) and 2020 (n = 3). Colours represent FOD encountered by seals, identified from in-situ temperature and salinity profiles recorded by CTD tags. Black lines indicate the mean position of the following oceanic fronts: South-Antarctic Front (SAF), Polar Front (PF), South Antarctic Circumpolar Current Front (SACCF) and Southern Boundary of the Antarctic Circumpolar Current (SBdy) extracted from Kim et al. (2014). (B) Average temperature and salinity profiles of the four FOD encountered by seals during their foraging trips. For each profile, the envelopes are bounded by the first and last quartiles and thus contain 50% of the profiles.

**Table 2:** Proportion of dives performed in each FOD visited by seven females SES from Kerguelen Islands equipped with high-resolution sonar and movement tags between 2018 and 2020. FOD were identified from in-situ temperature and salinity profiles recorded by CTD tags.

| Seal ID | FOD1 | FOD2 | FOD3 | FOD4 |
| --- | --- | --- | --- | --- |
| ml18_292a | 82 | 18 | 0 | 0 |
| ml18_293a | NA | NA | NA | NA |
| ml18_294a | 8 | 10 | 49 | 33 |
| ml19_295a | 36 | 63 | 0 | 1 |
| ml19_295c | 41 | 43 | 3 | 13 |
| ml19_296a | NA | NA | NA | NA |
| ml20_295a | 13 | 61 | 25 | 2 |
| ml20_296a | 9 | 52 | 33 | 6 |
| ml20_296c | 76 | 23 | 1 | 0 |

Bouts performed in FOD 4 were significantly deeper than those performed in FOD 1, 2 and 3 during both day and night (depth _day_ = 296 ± 169 m, 244 ± 165 m, 414 ± 159 m, 508 ± 138 m in FOD 1, 2, 3 and 4 respectively, depth _night_ = 224 ± 105 m, 179 ± 98 m, 287 ± 100 m, 254 ± 94 m in FOD 1, 2, 3 and 4 respectively, LMM, P < 0.001, Figure 3). Moreover, difference in day and night bout depths was more pronounced in FOD 3 and 4 (bout depth _day_ – bout depth _night_ = 99 ± 113 m and 205 ± 98 m respectively) compared to FOD 1 and 2 (bout depth _day_ – bout depth _night_ = 60 ± 90 and 42 ± 91 m respectively, LMM, P < 0.005).

**Figure 3:**
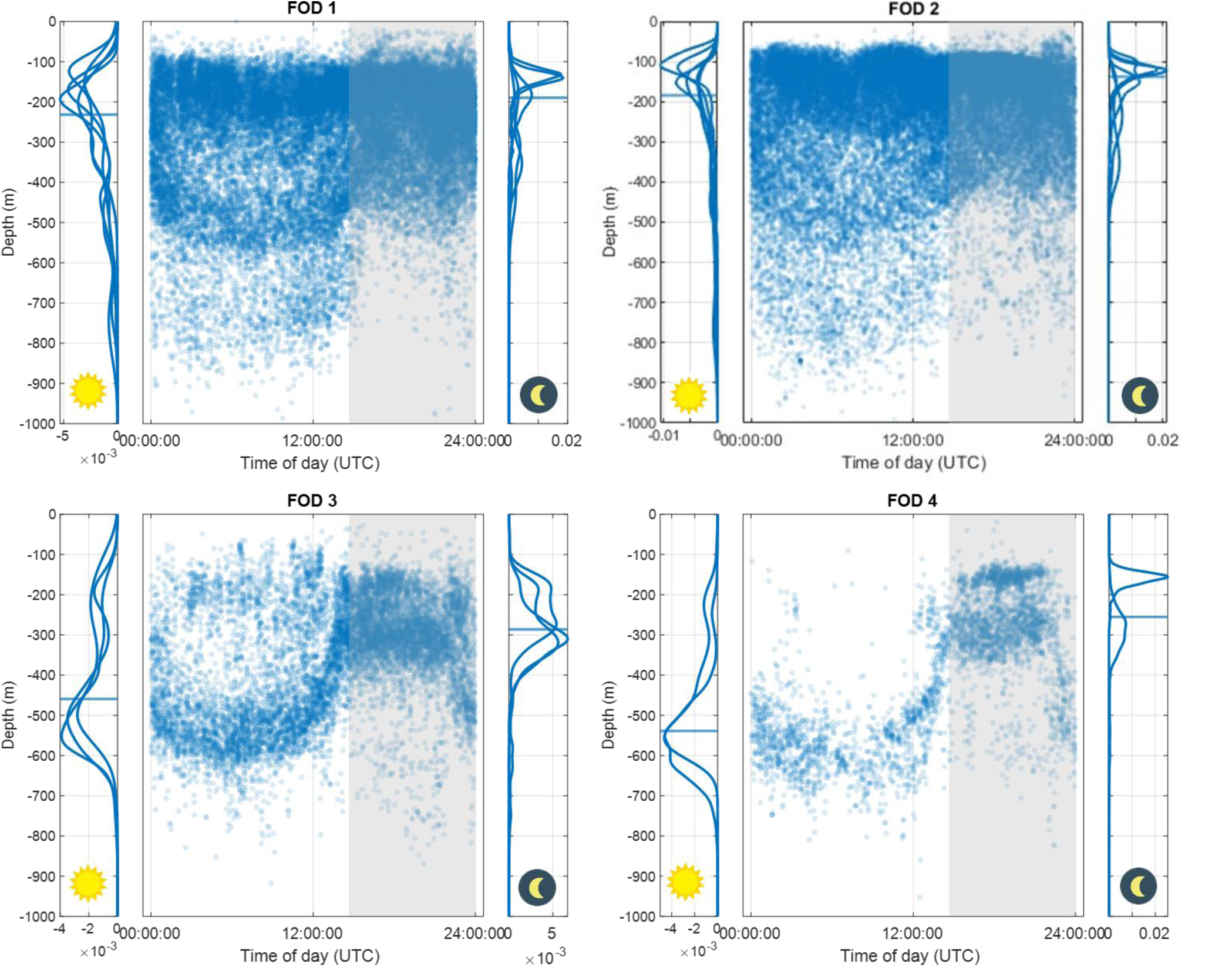
Depth of bouts over a 24h cycle recorded for the seven female SES equipped with sonar and movement tags in Kerguelen Islands from 2018 and 2020, separated by FOD visited. Each point represents a bout. The grey background represents nighttime. Curves on the left and right of each scatterplot represent the density distribution of the bout depth during day and night respectively reconstructed for each seal that spent more than 10% in the FOD, with the horizontal line representing the median depth.

The proportion of evasive prey was significantly higher in FOD 3 and 4 (93.9% and 90.6% respectively) than in FOD 1 and 2 (75.1 and 77.7% respectively) (GLMM, P < 0.001, Table 3). Similarly, prey tended to react at a larger distance in FOD 3 and 4 (72 ± 41 cm and 74 ± 44 cm respectively) than in FOD 1 and 2 (65 ± 38 cm and 56 ± 26 cm respectively) (LMM, P < 0.039, Table 3). The proportion of flashing prey was significantly higher in FOD 1 and 2 (28.5% and 27.9% respectively) than in FOD 3 (13.7%) (GLMM, P < 0.001). Acoustic sizes of prey encountered in FOD 4 were significantly greater than those of prey encountered in FOD 1, 2 and 3 (Kolmogorov-Smirnov test, P = 0.009, Table 3). Bouts performed in FOD 3 and 4 were significantly longer than those performed in FOD 1 and 2 (LMM, P < 0.021, Table 3).

**Table 3:**
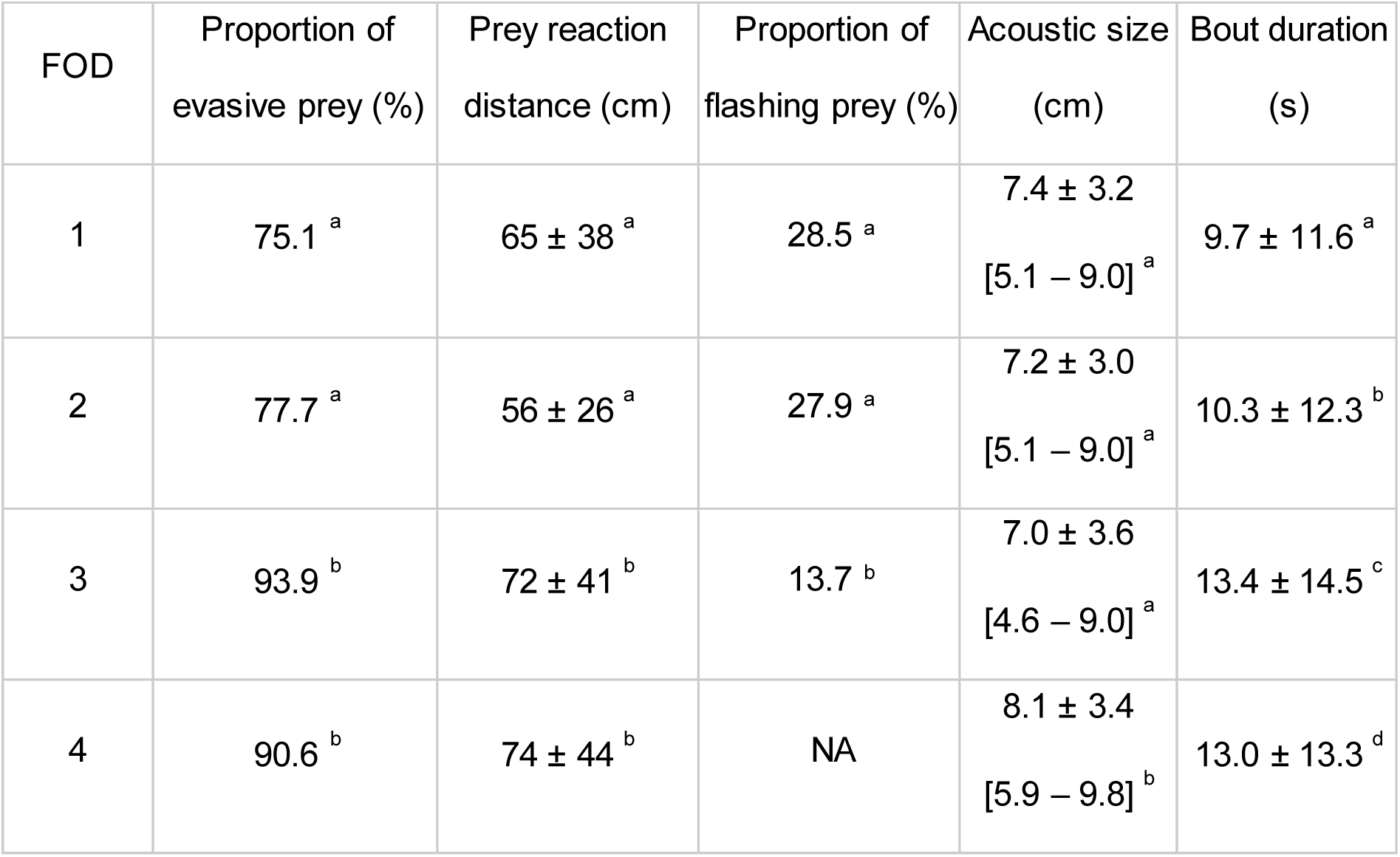
Bout duration, characteristics and behaviour of prey encountered by seven female SES equipped with high-resolution movement and sonar tags in Kerguelen Islands from 2018 to 2020, according to the FOD visited. Values are mean ± sd [1^st^ - 3^rd^ quartiles]. Letters in superscript indicate significant differences in prey characteristics between FOD (GLMM for proportion of evasive prey, LMM for prey reaction distance and bout duration, Kolmogorov-Smirnov test for acoustic size, P < 0.050).

### 3.4. Intra-functional oceanographic domain differences in prey characteristics and distribution

In FOD 1, 2 and 3, evasive prey were distributed into two distinct depth layers during the day, i.e. 100-300 m and 400-600 m, moving up to the sub-surface layers at night (Figure 4). In FOD 4, the sub-surface layer was not visible during the day and prey were mostly concentrated in the 500-700 m layer. On the opposite, non-evasive prey were mostly encountered in sub-surface layers, in depths ranging from 100 to 300 during both day and night, except in FOD 3 and 4. Overall, non-evasive prey were encountered significantly shallower than evasive ones during both day and night (LMM, P < 0.001, Figure 4). Similar to non-evasive prey, flashing prey were also mostly distributed in the sub-surface layers, i.e. 100-300 m during both day and night, except in FOD 3 and 4 where they were divided into two depth layers during the day, i.e. 100-300 m and 400-700 m (Figure 4). However, flashing prey were still encountered significantly shallower than non-flashing ones during both day (LMM, P = 0.017) and night (LMM, P = 0.010, Figure 4).

**Figure 4:**
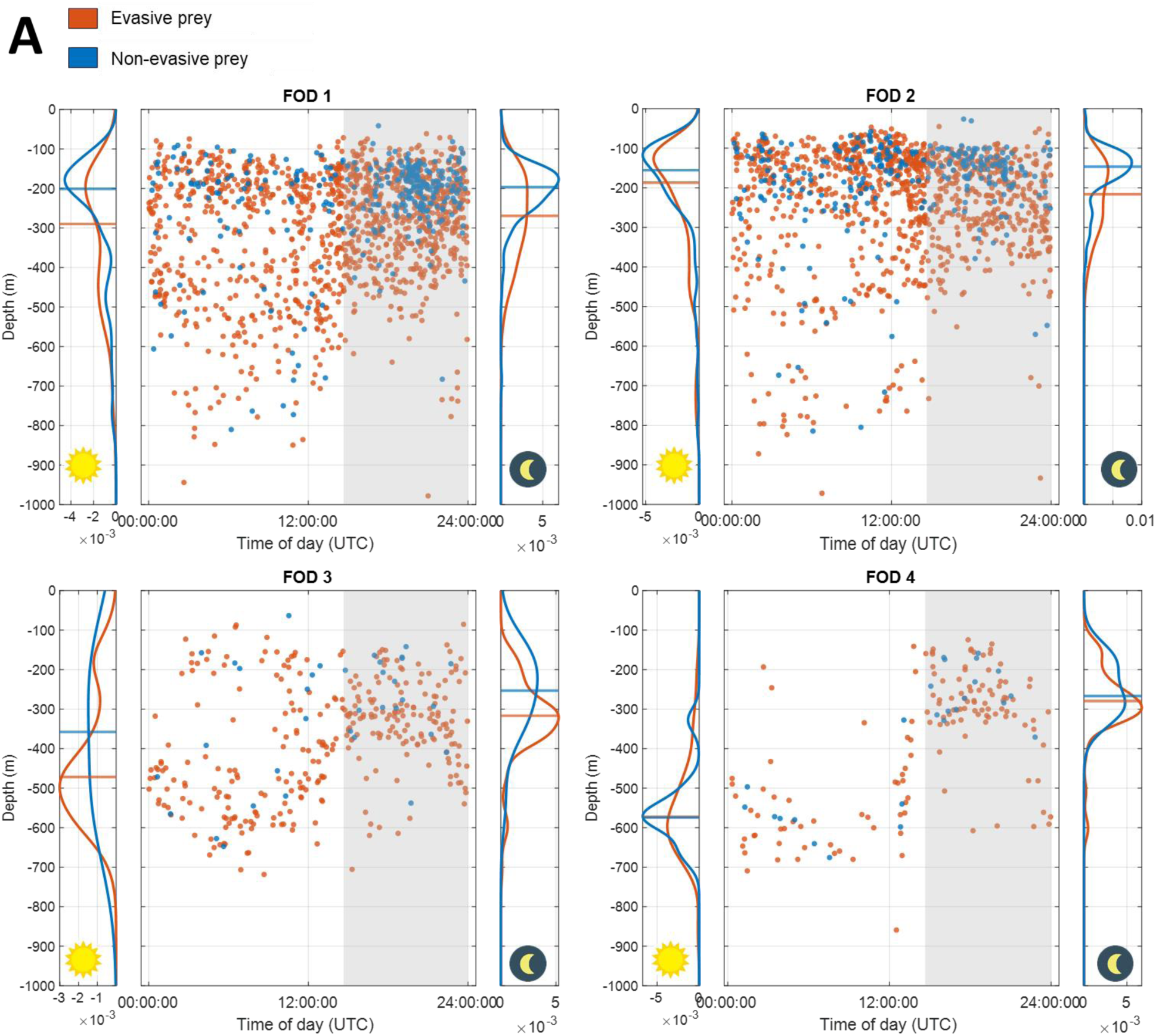
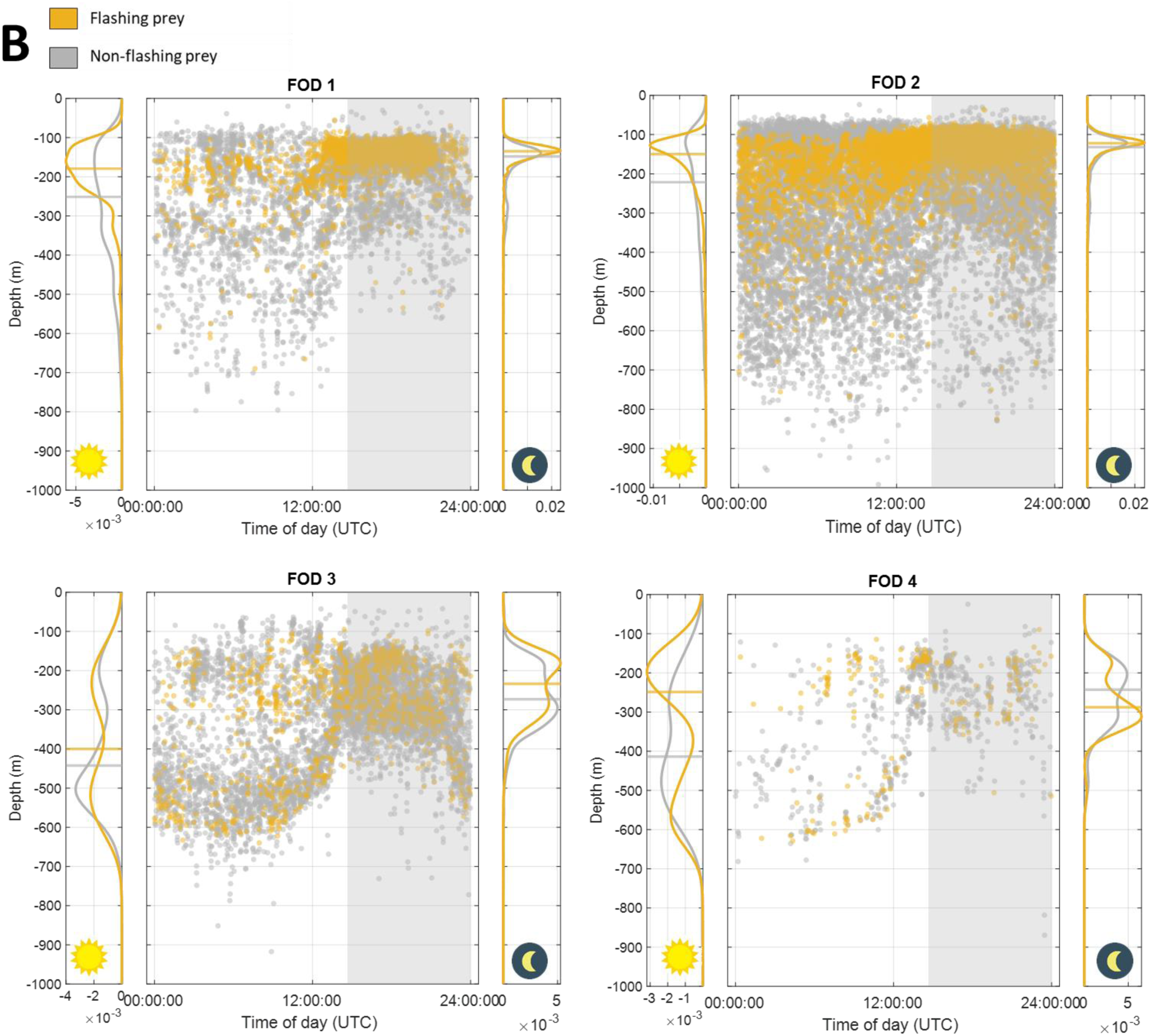
Depth of bouts over a 24h cycle recorded for female SES equipped with sonar and movement tags in Kerguelen Islands from 2018 and 2020. Each point represents a bout targeting (A) evasive or non-evasive prey (red and blue dots respectively, n = 6 seals), and (B) flashing or non-flashing prey (yellow and grey dots respectively, n = 2 seals). The grey background represents nighttime. Curves on the left and right of each scatterplots represent the density distribution of the bout depth during day and night respectively, with the horizontal line representing the median depth.

In all FOD, acoustic size of prey encountered in deeper waters was significantly greater than acoustic size of prey encountered in sub-surface layers (Kolmogorov-Smirnov test, P < 0.009, Figure 5).

**Figure 5:**
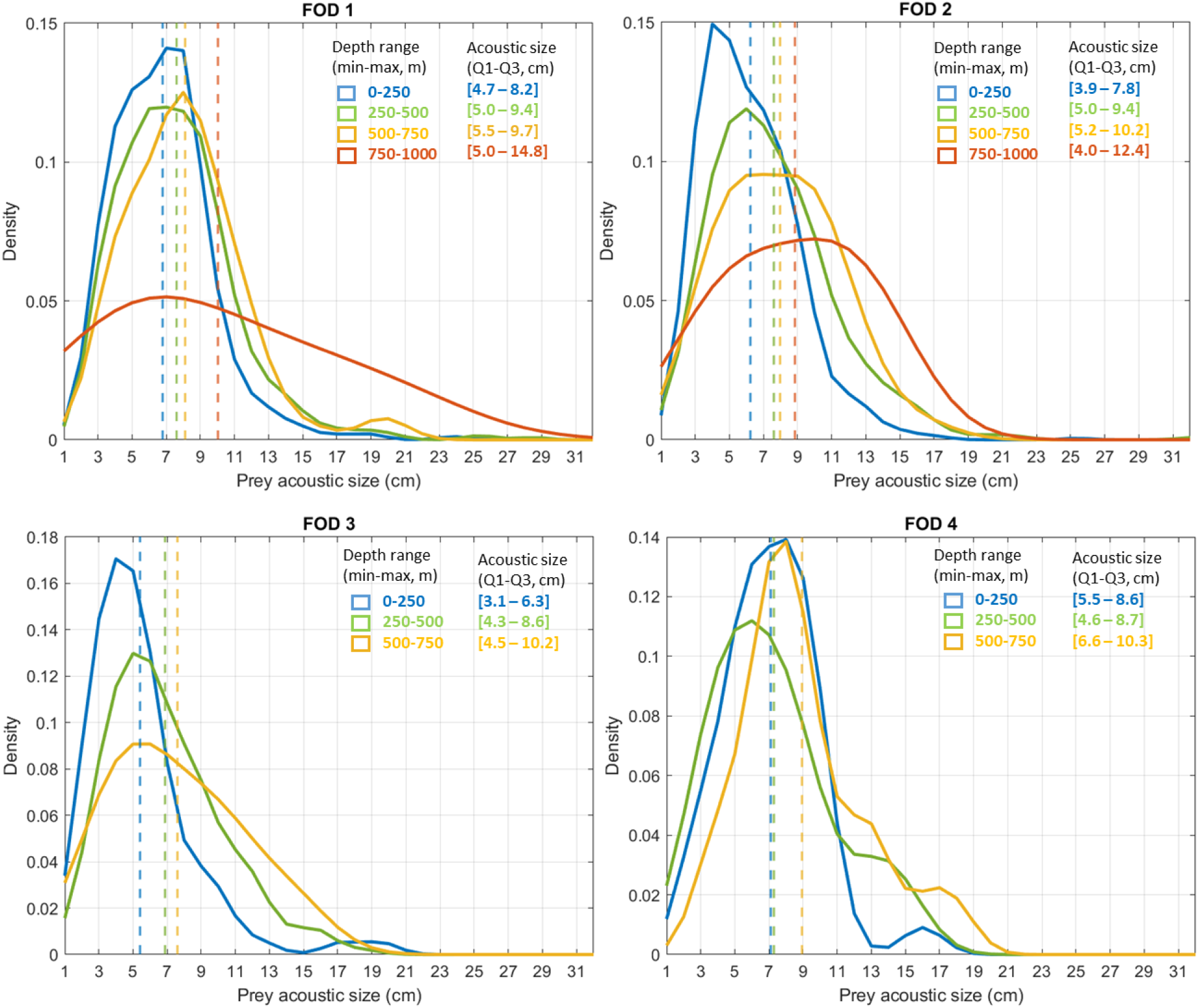
Prey acoustic size density distributions reconstructed for each depth class and for each FOD visited by seven female SES equipped with sonar and movement tags between 2018 and 2020 in Kerguelen Islands. The dashed lines represent the mean acoustic size calculated for each depth class.

In all FOD, bout duration increased with bout depth (LMM, P < 0.001): every 100 m, bout duration increased by 1.5 ± 0.04 s, 1.3 ± 0.04 s, 2.3 ± 0.09 s, and 1.0 ± 0.12 s for FOD 1, 2, 3 and 4 respectively.

Bout duration _FOD 1_ = (6.4 ± 0.6) + (0.015 ± 0.0004) · depth

Bout duration _FOD 2_ = (7.0 ± 0.7) + (0.013 ± 0.0004) · depth

Bout duration _FOD 3_ = (4.7 ± 0.8) + (0.023 ± 0.0009) · depth

Bout duration _FOD 4_ = (8.6 ± 1.2) + (0.010 ± 0.0012) · depth

## 4. Discussion

### 4.1. A new approach to infer mesopelagic prey distribution via deep-diving predators

We deployed on female SES an innovative miniaturised sonar tag that combines active acoustics with high-resolution movement and bioluminescence sensors to describe fine-scale characteristics of prey targeted by seals, i.e. their acoustic size, escape behaviour, bioluminescent behaviour and gregarious behaviour. Sonar tags were deployed simultaneously with CTD tags to record fine-scale in-situ oceanographic data during prey encounters, offering a unique opportunity to study prey characteristics and behaviour simultaneously in relation to oceanographic conditions in a Southern Ocean deep-diving predator. By deploying this tag on a deep-diving predator that forage on mesopelagic prey, the SES, we were able to get some insights into mesopelagic prey distribution and ecology, allowing a significant advance in our understanding of the functioning of the mesopelagic zone.

### 4.2. Insights into mesopelagic prey distribution and habitat

All prey were encountered in deep open waters outside the Kerguelen plateau, and were not associated with particular topographic features. Most prey encounters occurred in depths ranging from 100 to 800 m, and were characterised by a clear diel pattern with shallower encounters at night. However, this diel migration behaviour was most noticeable in warmer waters. Female SES are known to target mesopelagic fish such as myctophids (Cherel et al., 2008), whose ecology is consistent with our results (Catul et al., 2011). Prey acoustic size mostly ranged from 3 to 10 cm, which is close to the main size of myctophids inhabiting Kerguelen waters (Duhamel et al., 2000; Hulley and Duhamel, 2011), although it is important to remember that acoustic and actual sizes are not necessarily similar. Indeed, acoustic size is dependent on target orientation (Burwen et al., 2003), which cannot be controlled for here. However, when a prey escapes, it will likely be at some point oriented in the axis of the sonar beam, so measuring the maximum size of the trace of escaping prey should provide a reliable indication of the prey actual size. Therefore, data collected support the hypothesis that most targets insonified by the sonar tag during prey capture events had a body size consistent with myctophids, or more broadly small size mesopelagic fish and squids.

Even if data recorded by sonar tags do not allow for identifying species targeted by seals, it provided some precious information about their acoustic size, bioluminescent behaviour, escape behaviour and gregarious behaviour, allowing us to explore differences in prey types according to functional oceanographic domains (FOD) visited by seals. During their foraging trips, seals visited four FOD distributed along an increasing temperature gradient. FOD 1 and 2 were the coldest, with temperature below 2 and 2.5°C respectively. These two FOD might correspond to cold waters south of the Polar Front, characterised by an upper temperature of 2.8°C at 200 m (Park et al., 1993), with FOD 1 likely influenced by the southern boundary of Southern Antarctic Circumpolar Current, defined by a temperature of 1.6°C at 200 m (Park et al., 1993). On the opposite, FOD 3 and 4 were characterised by an upper temperature of 4-5°C and 7°C respectively and correspond to warmer water north to the Polar Front, and close to the Subantarctic Front for FOD 4 (Park et al., 1993).

Trawling and acoustic surveys around Kerguelen Islands showed contrasted mesopelagic communities between north and south of the Polar Front. *Krefftychthys anderssoni* (1-8 cm) and *Electrona antarctica* (3-10 cm) dominate the myctophid community south of the Polar Front, while *Electrona carlsbergi* (4-10 cm), *Gymnoscopelus fraseri* (5-11 cm)*, G. bolini* (15-28 cm) and *G. braueri* (4-14 cm) are more abundant north of the Polar Front (Collins et al., 2008; Duhamel et al., 2000; Hulley and Duhamel, 2011). The Polar Front is a transition zone between cold Antarctic waters and warmer subtropical waters (Gordon, 1971). By acting as a boundary between two different hydrological regions, it constitutes a barrier for many species, resulting in marked differences in marine communities north and south of the Polar Front (Koubbi, 1993). Therefore, we would expect to observe differences in prey behaviour and characteristics between FOD 1-2 (colder waters, south of the Polar Front) and FOD 3-4 (warmer waters, north of the Polar Front).

Prey were slightly larger in FOD 3-4 compared to FOD 1-2, however we expected the size difference to be more pronounced. Nevertheless, we noticed differences in prey behaviour depending on the FOD: the proportion of evasive prey was higher in warmer waters, while the proportion of flashing prey was lower. In addition, diel vertical migration patterns were more pronounced in warmer waters, suggesting that prey encountered north and south of the Polar Front are of distinct species, differing in terms of reactivity, bioluminescent behaviour and diel vertical migration behaviour. Acoustic data recorded by bioluminescence-sonar tags therefore provide novel insights on differences on mesopelagic prey behaviour according to oceanographic conditions.

Moreover, for each given FOD, we observed strong differences in prey acoustic size and behaviour according to depth: prey encountered deeper were larger than those encountered in sub-surface layers no matter the FOD visited. Therefore, prey size and behaviour seemed to be structured mostly vertically, as observed in previous studies showing that depth is one of the most important factors in the variability of mesopelagic communities (Cook et al., 2013; Kenchington et al., 2020; Sutton et al., 2010). Hulley et al. (2011) reported that within a myctophid species, larger and older individuals are usually encountered deeper than smaller ones. Therefore, vertical size structuration observed in all FOD visited by our tagged seals is consistent with previous findings and is likely to be related with differences in maturity stages between individual prey and not necessarily be due to differences in targeted species.

In addition to observing differences in prey size with depth, we also highlighted differences in prey behaviour along the water column. Bioluminescent prey were mostly encountered in sub-surface layers during both day and night, while we expected to observe them in deeper waters (Haddock et al., 2010). Moreover, non-evasive prey were concentrated to shallow waters during both day and night, while evasive prey were also encountered in deeper waters. This pattern was observed in all FOD visited by seals, even if it was more noticeable in colder FOD as few prey were encountered in sub-surface layers in warmer FOD. Although we would have expected that prey encountered in deeper and therefore colder waters were less evasive due to a slower metabolism (Torres and Somero, 1988), we assumed that deep prey, being larger, were more vigilant than smaller/younger individuals encountered in shallower waters due to an increasing predation risk with prey size (Møller and Erritzøe, 2014).

To summarise, prey encountered in colder FOD could mostly belong to Antarctic species such as *Electrona antarctica* or *Krefftichthys anderssoni,* with either adults *K. anderssoni* or juveniles *E. antarctica* in sub-surface layers during both day and night, and adult *E. antarctica* exhibiting diel migration being deeper waters during the day, and going back to the sub-surface layers at night (Duhamel et al., 2000). However, the fact that the sub-surface layers seemed dominated by small, non-evasive and bioluminescent prey that did not perform diel vertical migration may suggest that these prey could be other organisms than myctophids such as small bioluminescent gelatinous low energy content organisms or euphausids (Cotté et al., 2022; Trebilco et al., 2019). In warmer FOD, prey encountered could belong to species associated with the subtropical waters such as the genus *Gymnoscopelus,* larger and faster swimming more evasive myctophid species, usually encountered in deeper waters and known to perform diel vertical migration (Duhamel et al., 2000).

### 4.3. Where to hunt?

Seals concentrated most of their foraging activity in FOD 1 and 2, i.e. south of the Polar Front. As observed by Field et al. (2004), Guinet et al. (2014) and McMahon et al. (2019), the southern Polar Frontal zone appears to be a preferred area for seals, as they performed shallower dives in that area compared to FOD 3 and 4. The Polar Front is the main foraging area of many species of birds and marine mammals. This area is characterised by intense mesoscale activity, allowing the concentration of organic material and therefore a high biological production. In particular, the southern Polar Frontal zone is known to gather 80% of the myctophid biomass of the Southern Ocean (Lubimova et al., 1987), making this area particularly interesting for SES.

Our main hypothesis was that some habitats showed higher prey accessibility and/or prey quality and were therefore preferred by seals. We found that prey encounters were significantly deeper in warmer FOD during both day and night. This result is consistent with observations by Guinet et al. (2014), Biuw et al. (2007) and McIntyre et al. (2011), showing that bout depth increases with water temperature. Data recorded during trawling and acoustic surveys revealed a deepening of myctophid communi ty when getting closer to subtropical waters (Hulley and Duhamel, 2011), suggesting that prey were less accessible in warmer FOD compared to cold FOD. Moreover, in warmer FOD, prey tended to be slightly more evasive with higher flight initiation distances. A previous study made on the same dataset showed that seals capturing evasive prey were more likely to initiate energy-costing active chase, while non-evasive prey could be passively captured (Chevallay et al., in revision). Indeed, we also observed that bout duration, a proxy for prey capture difficulty (Goulet et al., 2020), was longer in warmer FOD, suggesting that prey captured in warmer waters induced longer handling and/or pursuit times. Goulet et al. (submitted) revealed that evasive prey were associated with a slower improvement in seals’ body condition compared to non-evasive prey by relating body condition gain to prey behaviour in SES. To our knowledge, the only studies on the energy content of mesopelagic prey around Kerguelen Islands focus on two or three myctophid species, and have not shown strong differences in energy content between the species studied (Lea et al., 2002; Van de Putte et al., 2006). So there likely is a trade-off between prey size (and energy content) and energy expenditure SES spent to capture them. More species should be studied to make assumptions on the nutritional value of the various prey encountered by elephant seals in the different FOD visited. However, differences in prey behaviour and accessibility between FOD suggest that larger and deeper prey encountered north of the Polar Front might be less accessible and more difficult to capture, increasing locomotion costs, which might make these environments less profitable for elephant seals.

### 4.4. Limits of the study

This study highlights the usefulness of the bioluminescence-sonar tag to infer mesopelagic prey distribution and habitat when deployed on deep-diving predators. However, the main limitation of this method is that sampling is biased by the foraging behaviour of the equipped predator. In particular, it might be difficult to disentangled inter-FOD differences with individual preferences, as not all seals visit all FOD. However, when looking for each individual the differences in foraging behaviour between FOD, inter-FOD differences were more pronounced than the inter-individual differences inside each FOD (see Figure 3), suggesting that there are actual structural differences between FOD that are not related to individual preferences. In addition, it is known that seals oversample areas that favour prey aggregation such as frontal systems (Bailleul et al., 2007; Dragon et al., 2010; Siegelman et al., 2019) and do not sample deep waters at night because of the diel migrating pattern of their prey (Slip et al., 1994). However, the small size of the tag makes it possible to deploy on other diving predator species such as fur seals or penguins that also feed on mesopelagic prey but in different vertical/horizontal areas (Bost et al., 2002; Boyd et al., 1991; Charrassin and Bost, 2001) to increase spatio-temporal covering and get a more complete picture of the distribution of mesopelagic prey in the Southern Ocean. Moreover, there is still some uncertainties regarding the nature of targets insonified by the sonar, as acoustic data can be very challenging to interpret. Subsequently, integrating a camera triggered by acoustic detection to the sonar tag could help validating the actual nature of acoustic targets, enabling to be more precise in the description of mesopelagic prey characteristics.

### 4.5. Conclusion

Using a novel multi-sensor approach deployed on a deep-diving predator, the SES, we were able to explore the functional relationships between oceanographic parameters, distribution and ecology of mesopelagic prey, and foraging behaviour of their predators. We highlighted differences in prey size and behaviour with depth inside each FOD, as well as differences in prey accessibility and behaviour between FOD. Climate change leads to a deep-reaching ocean warming, and this warming is faster in the Southern Ocean compared to the global average (Gille, 2008). Increase in water temperature can considerably alter organism distributions, e.g. studies made on several fish stocks (Nye et al. (2009) on Northeast USA stocks; Dulvy et al. (2008) on North Sea stocks) revealed a deepening of fish stocks with increasing temperature to reach colder waters that are closer to their temperature ranges (Jorda et al., 2020). Therefore, with ocean warming, prey might be increasingly less accessible, forcing seals to dive deeper, which could lead to significant increase in foraging energy costs and exercise strong selective pressure on larger animals, which are found to dive deeper (Piot et al., 2023).

## Author contributions

MC, TJDD, CG: conceptualization

MC, TJDD, PG, NF, CG, BP: methodology

MC, CC: formal analysis

MC, PG, BP: data curation

MC: writing – original draft

MC, TJDD, PG, NF, CC, BP, CG: writing – review and editing

TJDD, CG: supervision

## Acknowledgments

SES data were gathered as part of the “Système National d’Observation: Mammifères Echantillonneurs du Milieu Océanique” (SNO-MEMO, PI. C. Guinet) and in the framework of the ANR HYPO2. Fieldwork in Kerguelen was supported by the French Polar Institute (Institut Polaire Français Paul Emile Victor) as part of the CyclEleph programme (n. 1201, PI C. Gilbert) and with the help of the Ornitho-eco programme (n. 109, PI C. Barbraud). Data acquisition was also supported by CNES-TOSCA (Centre National d’Études Spatiales). The authors wish to thank all individuals who, over the years, have contributed to the fieldwork of deploying and recovering tags at Kerguelen Island. We also thanks Yves Cherel for his help in interpreting the data, and Mark Johnson for providing tags, software and codes for data analysis.

